# A computational observer model of spatial contrast sensitivity: Effects of photocurrent encoding, fixational eye movements and inference engine

**DOI:** 10.1101/759811

**Authors:** Nicolas P. Cottaris, Brian A. Wandell, Fred Rieke, David H. Brainard

**Affiliations:** Department of Psychology, University of Pennsylvania, Philadelphia, PA, USA; Department of Psychology, Stanford University, Stanford, CA, USA; Department of Physiology & Biophysics, University of Washington, WA, USA

**Keywords:** contrast sensitivity function, computational modeling, fixational eye movements, photocurrent, phototransduction, spatial pooling, inference engine

## Abstract

We have recently shown that using the information carried by the mosaic of cone excitations of a stationary retina, the relative spatial contrast sensitivity function (CSF) of a computational observer has the same shape as a typical human subject. Absolute human sensitivity, however, is lower than the computational observer by a factor of 5 to 10. Here we model how additional known features of early vision affect spatial contrast sensitivity: fixational eye movements and the conversion of cone photopigment excitations to cone photocurrent responses. For a computational observer that uses a linear classifier applied to the responses of a stimulus-matched linear filter, fixational eye movements substantially change the shape of the spatial CSF, primarily by reducing sensitivity at spatial frequencies above 10 c/deg. For a computational observer that uses a translation-invariant calculation, in which decisions are based on the squared response of a quadrature-pair of linear filters, the CSF shape is little changed by eye movements, but there is a two-fold reduction in sensitivity. The noise and response dynamics of conversion of cone excitations into photocurrent introduce an additional two-fold sensitivity decrease. Hence, the combined effects of fixational eye movements and phototransduction bring the absolute sensitivity of the translation-invariant computational observer CSF to within a factor of 1 to 2 of the human CSF. We note that the human CSF depends on processing of the initial representation by many thalamic and cortical neurons, which are individually quite noisy. Our computational modeling suggests that the net effect of this noise on contrast-detection performance, when considered at the neural population level and behavioral level, is quite small: the inference mechanisms that determine the CSF, presumably in cortex, make efficient use of the information available from the cone photocurrents of the fixating eye.

## Introduction

The spatial contrast sensitivity function (CSF) is a fundamental characterization of human vision: It specifies the amount of contrast required for a visual system to detect sinusoidal contrast modulation at different spatial frequencies. The human CSF is single-peaked, increasing with spatial frequency to about 3-5 c/deg (cycles per degree) and then declining steadily until the resolution limit, near 60 c/deg (Robson, 1966; Campbell & Robson, 1968; Kelly, 1977). The shape of the falling limb of the human CSF is parallel to that of an ideal observer that makes optimal use of the information carried by the excitations of the cone photoreceptors in a model foveal retinal mosaic (Banks, Geisler, & Bennett, 1987). This alignment indicates that blurring by the eye’s optics and cone apertures play an important role in limiting human contrast sensitivity. The absolute sensitivity of the ideal observer CSF, however, greatly exceeds that of human observers. This leads to the question of what visual mechanisms, not included in the ideal observer calculations, account for the lower sensitivity.

The ideal observer uses a decision rule that requires exact knowledge of the visual stimulus. Our recent work (Cottaris, Jiang, Ding, Wandell, & Brainard, 2019) relaxes this assumption by modeling an observer that learns the decision rule from labelled stimulus-response data. Indeed, it is to highlight this difference that we use the term *computational observer* (Farrell, Jiang, Winawer, Brainard, & Wandell, 2014). Using open-source and freely available software (ISETBio), we confirmed the essential features of the classic ideal observer results. The software extends the ideal observer work, allowing us to explore how variation in optics, cone mosaic structure and choice of learned decision model (*inference engine*) affect the spatial CSF (Cottaris et al., 2019). Accounting for these factors, which affect the encoding of every visual stimulus, did not change the shape of the falling limb of the computational observer CSF. There remained, however, a 5 to 10-fold difference in absolute sensitivity between the computational and human CSFs.

The present paper extends the analysis further into the visual system. We incorporate two additional factors into the ISETBio simulations that influence the visual encoding of every stimulus: (a) spatial uncertainty introduced by fixational eye movements and (b) sensitivity regulation and noise introduced by the conversion of cone excitations into cone photocurrent. Fixational eye movements, which include slow drifts and microsaccades, translate the retinal image with respect to the cone mosaic and introduce spatial stimulus uncertainty. Drifts translate the retinal image along Brownian motion-like curved paths which change direction frequently, with instantaneous mean velocities in the range of 30 – 90 arc min/sec (Cherici, Kuang, Poletti, & Rucci, 2012). Microsaccades occur between periods of drift, every 500-2000 msec, and induce very fast retinal image translations with speeds in the range of 4 – 100 deg/sec (Martinez-Conde, Macknik, Troncoso, & Hubel, 2009). Inference mechanisms in the visual system must confront this uncertainty. Although our model of fixational eye movements includes the ability to model both drift and microsaccades, here we only examine the effect of drift, as the modeled stimulus duration is sufficiently short (100 msec) that microsaccades rarely occur.

The conversion of cone excitations into photocurrent (phototransduction) introduces nonlinear amplification and compression of the cone excitation signal (Endeman & Kamermans, 2010) and additive noise that is largely stimulus-independent (Angueyra & Rieke, 2013). As mean light level increases, the effect of these two factors exceeds the uncertainty caused by Poisson noise in the cone excitations. Additional effects are also introduced by phototransduction, such as background-dependent changes in the temporal dynamics of the cone response and response asymmetries between increments and decrements (Endeman & Kamermans, 2010; Angueyra, 2014).

To foreshadow our main result, addition of fixational drift and phototransduction into the analysis pipeline closes much of the overall gap between computational and human CSFs, while mostly retaining the agreement in the shape of sensitivity falloff as spatial frequency increases. We say mostly, because our simulations reveal that the combined effect of fixational eye movement and phototransduction introduces an additional, small attenuation factor, which is spatial frequency - dependent. Thus, although the CSF of a human subject depends on processing by many thalamic and cortical neurons, which are individually quite noisy, our computational modeling suggests that the net effect of this noise on contrast detection, when considered at the neural population level, is quite small: the inference mechanisms that determine the CSF, presumably in cortex, make efficient use of the information available from the cone photocurrents of the fixating eye.

## Overview of Computational Model of Early Vision

Evaluating the significance of a wide array of visual system factors requires computational modeling. To meet this challenge we are developing ISETBio (Farrell et al., 2014; Jiang et al., 2017; Cottaris et al., 2019) and related software packages (Lian, MacKenzie, Brainard, Cottaris, & Wandell, 2019) as open-source software resources (https://github.com/isetbio/isetbio, https://github.com/isetbio/ISETBioCSF, https://github.com/ISET/iset3d). The software enables specification of visual scene radiance (including both three-dimensional scenes and stimuli presented on planar displays), modeling the transformation of scene radiance through the eye’s optics to retinal image formation, calculation of cone excitations in the retinal cone mosaic, modeling of phototransduction within the cones, simulation of fixational eye movements, and implementation of inference engines for relating visual representations to performance on psychophysical tasks. The computational pipeline for scenes represented on planar displays through to the level of cone photopigment excitations is described in detail elsewhere (Cottaris et al., 2019). The work here describes extensions to include fixational eye movements, which translate the retinal image over the cone mosaic, and a model of phototransduction which converts quantal cone excitation events into current flow through the cone outer segment membrane.

An overview of the computational pipeline is shown in Figure 1. The retinal image^1^ is the spectral irradiance incident at the retina. The irradiance is computed from the spectral radiance emitted by a screen using a model of the eye’s optics (Cottaris et al., 2019). This part of the pipeline accounts for critical physiological optics factors such as pupil size, wavelength-dependent transmission through the crystalline lens and on-axis wave-front aberrations of the eye’s optics. Fixational eye movements generate a temporal sequence of retinal images, two frames of which are depicted in Figure 1B. The cone mosaic samples the retinal image sequence resulting in a sequence of cone mosaic excitations; two frames are shown in the left panels of Figure 1C. The entire sequence of cone excitations over the simulated 150 msec interval is depicted for a single cone in the right panel of Figure 1C. The blue line depicts the noise-free response and the black line depicts a single noisy response. The sequence of cone excitations drives the cone’s photocurrent noise-free response, which is computed via a biophysically-based phototransduction model (Figure 1D). Photocurrent noise, whose spectral density is matched to that of photocurrent responses in primate cones, is added to the noise-free response to produce a photocurrent response instance, depicted in black in Figure 1D.

**Figure 1:**
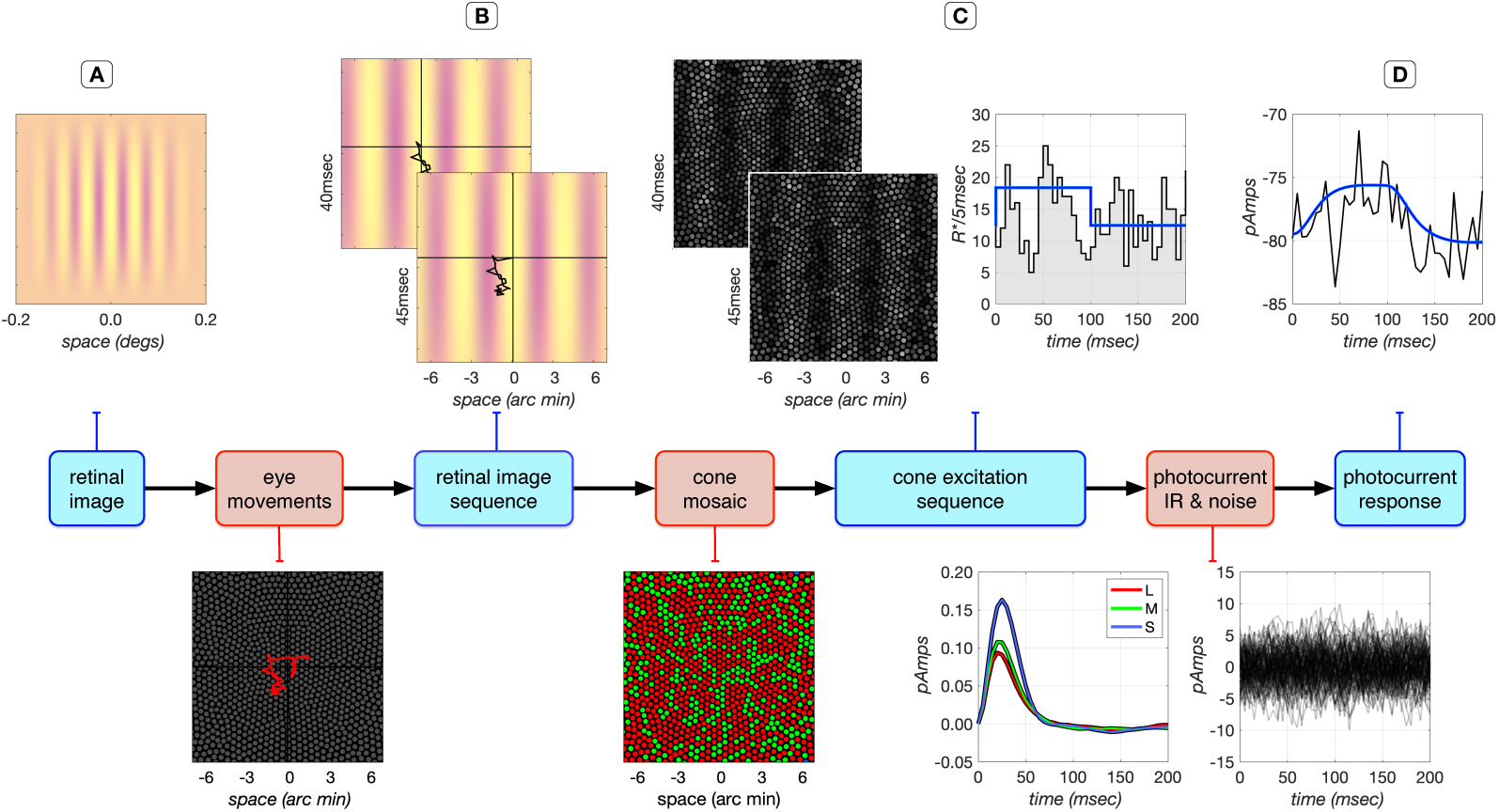
ISETbio flowchart for computation of photocurrent responses in the presence of fixational eye movements. **A**. Retinal image of a 100% contrast, 16 c/deg achromatic grating stimulus pulsed for 100 ms. The yellow tint of the rendered image is due to short wavelength absorption by the lens. Fixational eye movements, depicted as red traces over the central 14 arc min of the cone mosaic in the bottom plot, translate the retinal image, creating a temporal sequence of retinal images. **B**. Two frames of the retinal image sequence (central 14 arc min) at 40 and 45 msec after stimulus onset. Black lines trace the retinal image translation up to the time of the frame, and the crosshairs show the center of the translating retinal image on the display. The retinal image sequence is sampled by the cone mosaic, generating a sequence of cone excitations. **C**. Two frames of the cone excitation sequence at 40 and 45 msec are depicted on the left, and the time course of excitations for a single cone is plotted on the right. The mean cone excitation response in the absence of eye movements is depicted in blue and a single, noisy cone excitation response instance in the presence of fixational eye movements is depicted in black. The noisy instance follows a Poisson distribution with a time-varying rate parameter equal to the expected number of cone excitation events in each time bin. **D**. The noise-free excitation sequence of each cone is convolved with the photocurrent impulse response to generate the noise-free photocurrent response (blue) and noisy photocurrent response instances (black). For a given mean cone excitation rate, the mean photocurrent response is approximated using a temporal linear filter, whose impulse response is computed separately for each cone type, using a biophysically-based model of phototransduction. The red, green and blue curves in the bottom left panel depict the L-, M-, and S-cone photocurrent impulse response functions for this mean. The S-cone impulse response has significantly higher gain than that of the L- and M-cones, because mean S-cone excitation is low which sets the photocurrent gain high. Photocurrent response instances are generated by adding stimulus-independent Gaussian noise (depicted in black at the right bottom panel) to the mean photocurrent response.

## Results

On each trial in a two-interval forced-choice spatial contrast sensitivity experiment, the subject is presented with two stimulus intervals, one of which contains a spatially-uniform pattern (null stimulus), and one which contains a sinusoidal grating pattern (test stimulus). The subject reports which interval contains the test. For each spatial frequency, contrast is varied across trials and percent correct is measured as a function of contrast. From such data, the contrast corresponding to a criterion percent correct is taken as detection threshold, and sensitivity is given by the reciprocal of threshold contrast.

Figure 2 depicts examples of cone mosaic excitation and photocurrent responses to a 16 c/deg, 100% contrast, 100 msec test stimulus, with a mean luminance of 34 cd/m^2^. The mosaic’s mean cone excitation response in the absence of fixational eye movements is shown in Figure 2A. The cone excitation responses increase with eccentricity because cone aperture increases with eccentricity. Figure 2B depicts a single noisy instance of a differential (test-null) spatiotemporal cone excitation response for cones along the horizontal meridian of the mosaic, again in the absence of fixational eye movements. A clear response modulation can be seen during the stimulus presentation duration (0-100 msec). Figure 2C depicts a single noisy instance of a differential (test-null) spatiotemporal photocurrent response. The stimulus-induced modulation in the photocurrent response is somewhat blurred over time and the modulation is more noisy than the cone excitation response. Figure 2D depicts four instances of fixational eye movement paths each computed for a period of 150 msec. Different eye movement trajectories start at random locations, but the trajectories are constrained so that their centroids are all at the origin. The spatiotemporal differential cone excitation response during one fixational eye movement instance (green line in Figure 2D) is depicted in Figure 2E. The response modulation is tilted in the space-time domain due to jitter in retinal image position along the horizontal axis. The corresponding photocurrent differential spatiotemporal response is depicted in Figure 2F. Note that the stimulus modulation is barely visible in this representation. Two factors contribute to this. First, the convolution of the jittered cone excitation response with the photocurrent impulse response smears the cone excitation response in time. Second, the photocurrent response signal to noise ratio (SNR) is significantly lower than the SNR of the cone excitation response, as can be seen by comparing Figures 2B and 2C.

**Figure 2:**
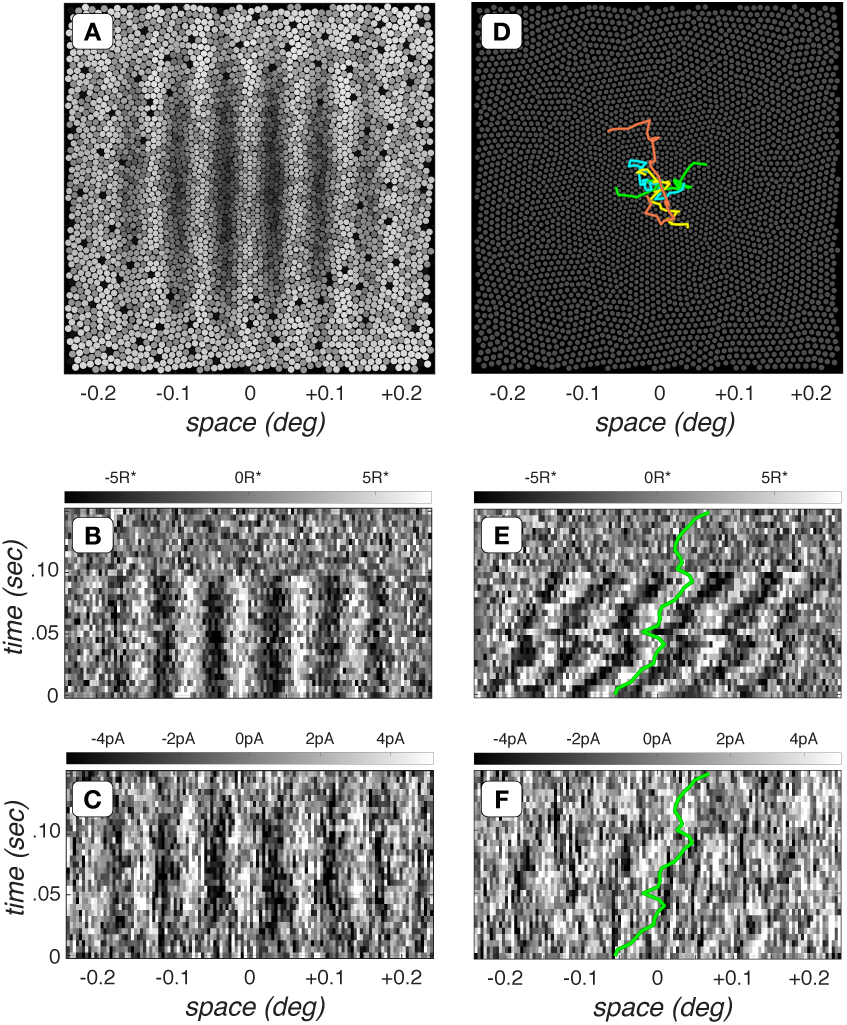
Spatiotemporal dynamics of cone mosaic excitation and photocurrent responses. **A.** Mosaic mean cone excitation response to a 16 c/deg, 100% contrast, 34 cd/m^2^ mean luminance grating, flashed for 100 msec. Excitation level is depicted by the gray scale value. S-cones are weakly excited and appear black. This is primarily due to selective absorption of short-wavelength light by the lens and macular pigment. **B.** Each column represents the differential (test - null) spatiotemporal cone excitations of a cone on the horizontal meridian (no fixational eye movements). Gray-scale color bar denotes excitation level in *R*^*^*/*cone*/*5 msec **C.** Differential photocurrent response of cones along the horizontal meridian. Gray-scale color bar denotes response level in pico Amperes (pA). **D.** Four instances of fixational eye movements, each computed for a period of 150 msec. **E.** Differential cone excitation response during one fixational eye movement instance (green trace in **C**). **F.** Differential photocurrent spatiotemporal response during the same fixational eye movement instance. The green lines in **E** and **F** show the horizontal position of a single cone.

### Impact of fixational eye movements

We begin our computational assessment of contrast sensitivity by examining the impact of fixational drift at the level of cone excitations. The gray disks in Figures 3A and 3B depict the contrast sensitivity function (CSF) in the absence of fixational eye movements. Performance is estimated using a computational observer which employs a linear support vector machine (SVM) classifier operating on the output of a stimulus-matched spatial pooling filter (template), which linearly sums cone responses over space at every time instant (Cottaris et al., 2019), and which we term the *SVM-Template-Linear* computational observer. This CSF serves as a baseline for assessing the impact of fixational eye movements.

**Figure 3:**
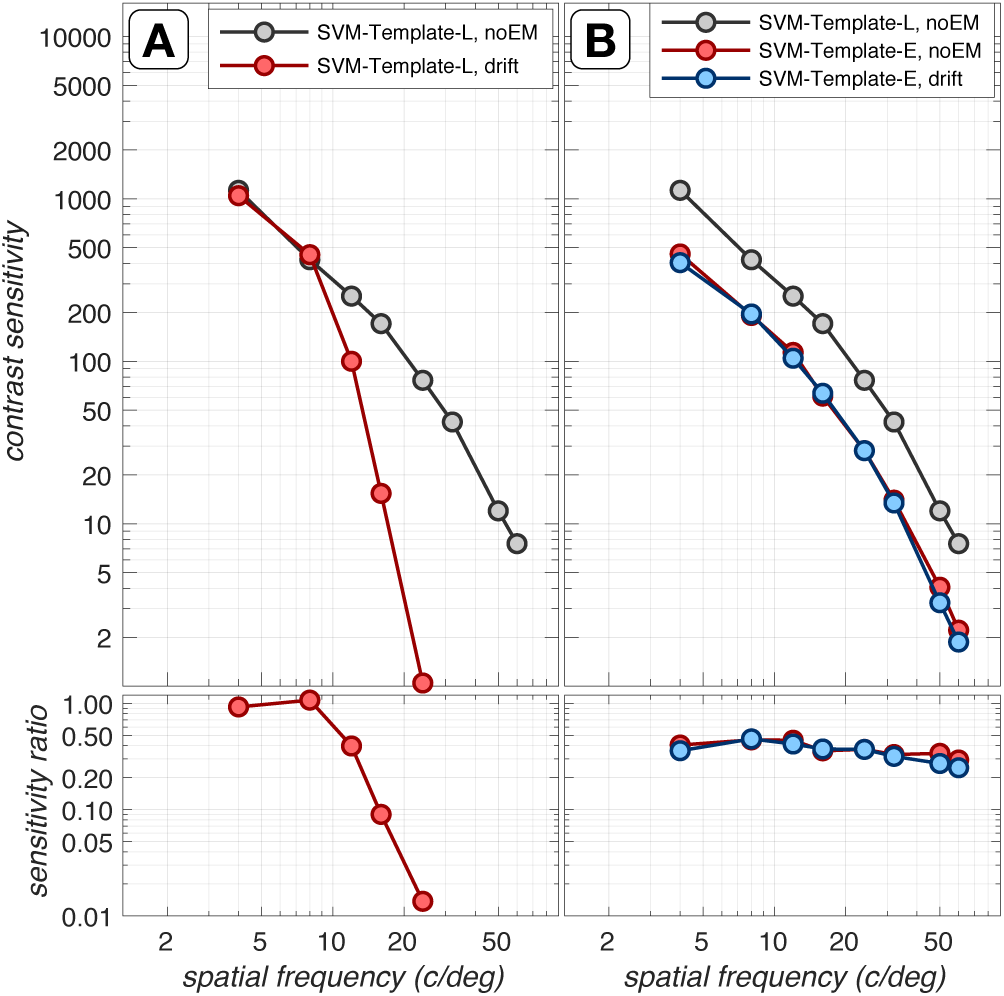
Impact of fixational eye movements. Top panels: Contrast sensitivity functions (CSF) at the level of cone excitations, computed for a 3 mm pupil, typical subject wavefront-based optics, and eccentricity-based cone mosaics (Cottaris et al., 2019). Bottom panels: Ratios of CSFs with respect to the reference CSF, which is computed using the SVM-Template-Linear observer in the absence of fixational drift (gray disks). **A**. CSFs computed using the SVM-Template based computational observer in the absence (gray disks) and presence of fixational drift (red disks). **B**: CSFs computed using the SVM-Template-Energy observer in the absence (red disks) and presence (blue disks) of fixational drift.

The CSF computed in the presence of drift fixational eye movements using the same SVM-Template-Linear observer is depicted by the red disks in Figure 3A. Note, the dramatic loss in sensitivity as spatial frequency exceeds 10 c/deg. Indeed, a threshold cannot be obtained beyond 24 c/deg. For this computational observer, fixational eye movements cause significant misalignment between the retinal image and the observer’s stimulus-matched filter. This decreases the SNR of the observer’s filter response in a spatial-frequency-dependent manner, leading to the rapid falloff in classifier performance with increasing spatial frequency.

A computational observer that is less susceptible to the effects of retinal image jitter can be constructed by employing a pair of stimulus-matched spatial pooling filters which have a spatial quadrature relationship (see Figure 7D,E), and whose outputs are squared (Cottaris et al., 2019; Greene, Gollisch, & Wachtler, 2016). We term this observer, the *SVM-Template-Linear* computational observer. This computation, often referred to as an energy computation, was introduced to explain retinal and cortical neuron responses that are independent of stimulus spatial phase within the neuron’s receptive field (Hochstein & Shapley, 1976; Emerson, Bergen, & Adelson, 1992; Ohzawa, DeAngelis, & Freeman, 1990). The SVM-Template-Energy computational observer applies a linear SVM classifier to the energy responses.

**Figure 4:**
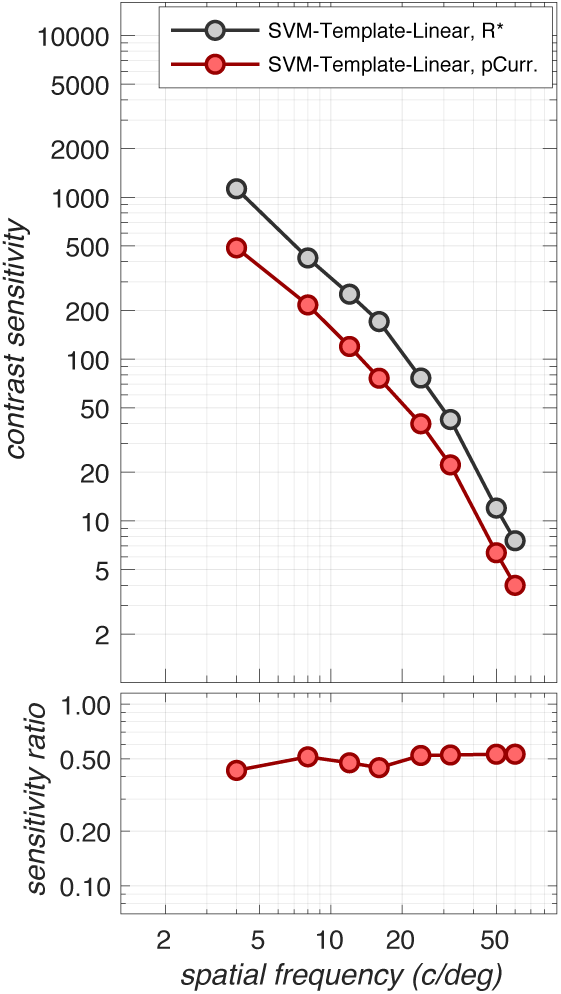
Impact of phototransduction. Contrast sensitivity functions computed at the level of cone excitations (gray disks) and at the level of photocurrents (red disks) in the absence of fixational eye movements. These were obtained using the SVM-Template-Linear computational observer, computed for a 3 mm pupil, typical subject wavefront-based optics, and eccentricity-based cone mosaics. The transformation from cone excitations to photocurrent results in a sensitivity loss of a factor of 2 to 2.5. Note that this sensitivity loss is specific to mean light level (here 34 cd/m^2^) and stimulus duration (here 100 msec). Stimuli presented at different adapting light levels and/or for different durations will be affected differently.

**Figure 5:**
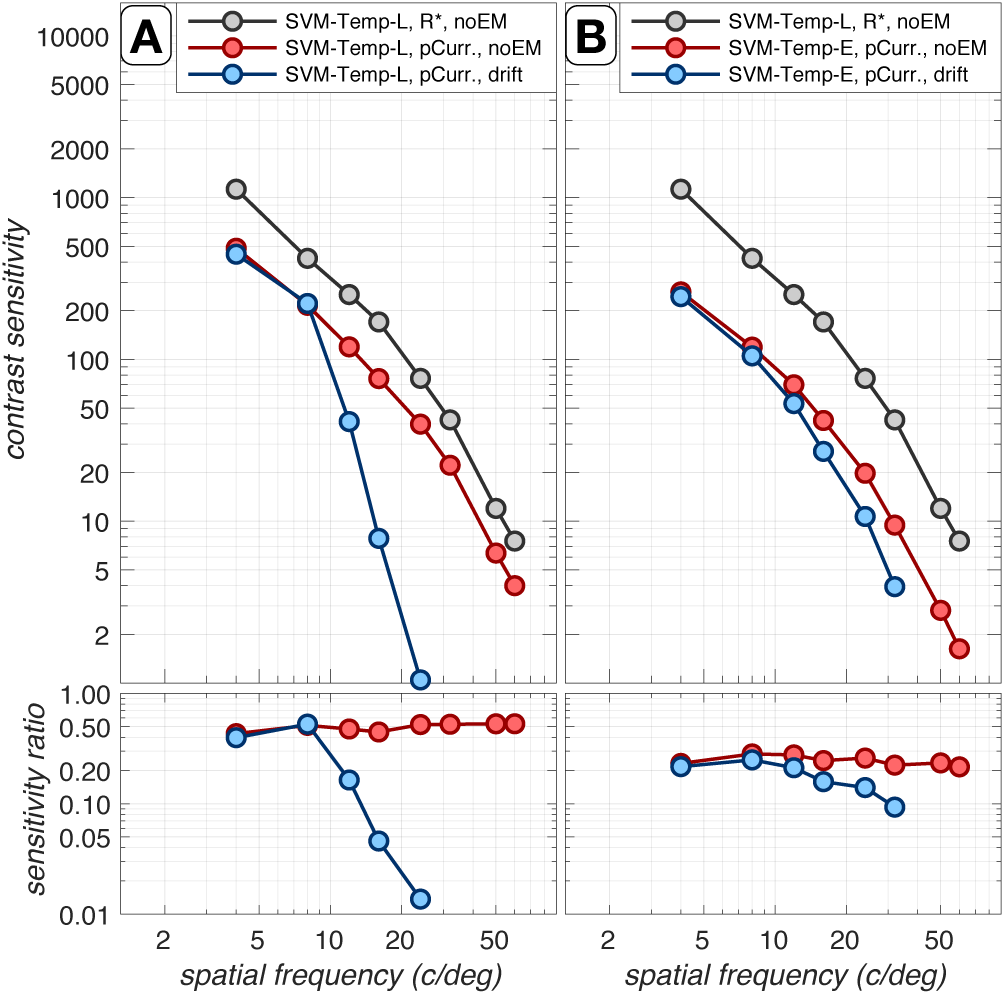
Combined effect of fixational eye movements and phototransduction. Contrast sensitivity functions for a 3 mm pupil, typical subject wavefront-based optics, and eccentricity-based cone mosaics. **A**. CSFs computed using the SVM-Template-Linear based computational observer. **B**: CSFs obtained with the SVM-Template-Energy computational observer. Gray disks: Reference CSF computed at the level of cone excitations in the absence of fixational eye movements. Red disks: CSFs computed at the level of photocurrent in the in the absence of fixational eye movements. In **A**, red disks are replotted from Figure 4. Blue disks: CSFs computed at the level of photocurrent in the in the presence of fixational eye movements.

**Figure 6:**
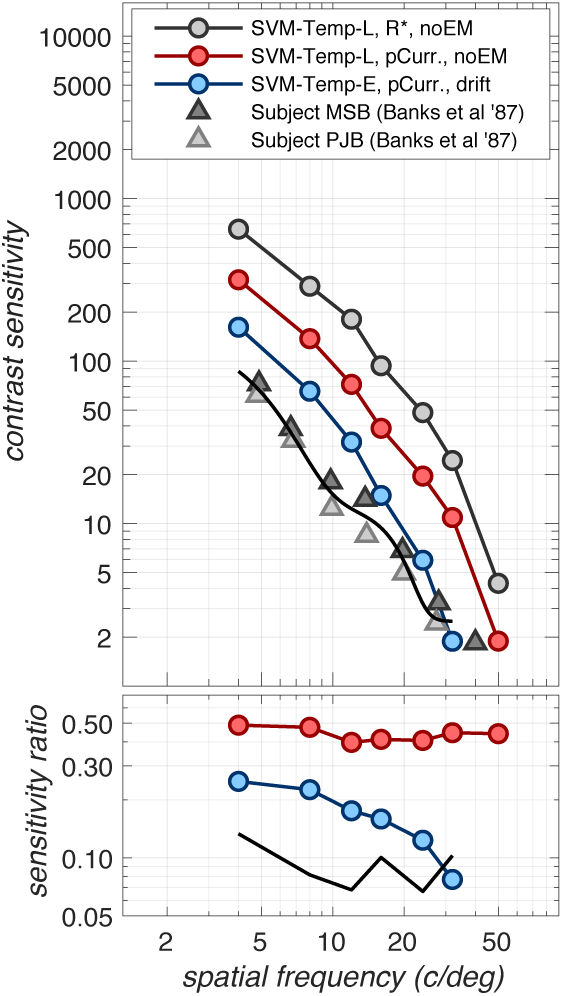
Comparing human and computational CSFs. Spatial CSFs are computed for 2 mm pupils, matching the psychophysical conditions of Banks et al. (1987). Gray disks depict the CSF at the level of cone excitations in the absence of fixational eye movements computed using the SVM-Template-Linear observer, which serves as the reference CSF. Red disks depict the CSF at the level of photocurrent also in the absence of eye movements and using the SVM-Template-Linear observer. Blue disks depict the CSF at the level of photocurrent in the presence of fixational eye movements using the SVM-Template-Energy observer. Triangles depict the CSFs measured in two subjects by Banks et al. (1987).

**Figure 7:**
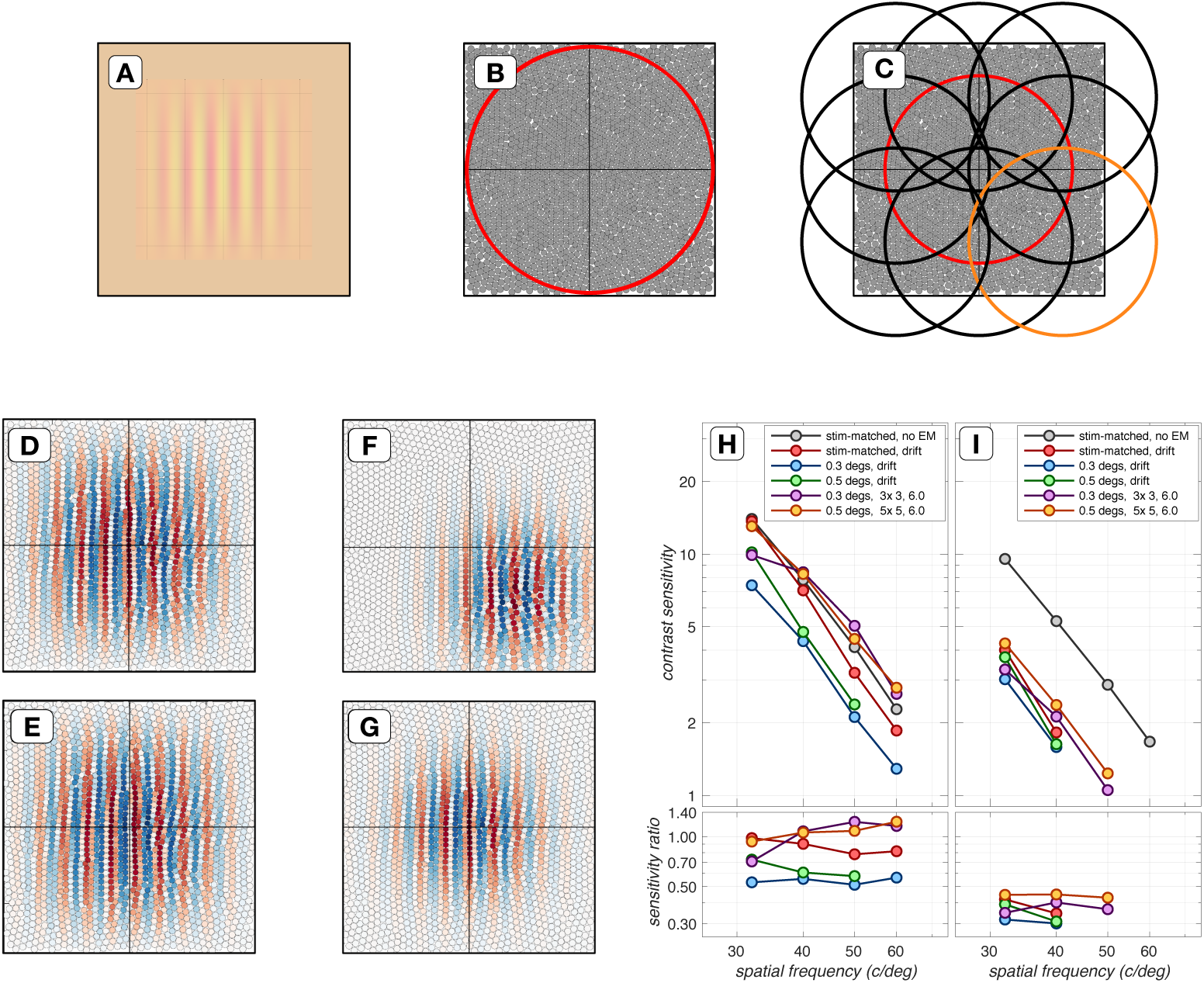
Impact of spatial pooling [Suppl. Fig. S0]. High spatial frequency regime of contrast sensitivity functions computed using different spatial pooling mechanisms. **A.** Retinal image of a test stimulus whose spatial support extends over a 0.25x *×* 0.25 deg region. **B.** The spatial pooling extent of a single energy mechanism integrating cone responses over a 0.33 *×* 0.33 deg region (red circle) is depicted superimposed on the cone mosaic. **C.** The spatial pooling extents of an ensemble of 9 spatially-offset energy mechanisms arranged in a 3 ×3 grid are depicted in black, red and orange circles. **D. and E.** The quadrature-phase pooling kernels for the single, spatially-extended energy mechanism depicted in the **B** panel. **F. & G.** Cos-phase pooling kernels for 2 filters in the 3 × 3 ensemble, outlined in red and orange in the **C** panel. **H. & I.** Performance of different spatial pooling mechanisms at the level of isomerizations and photocurrents, respectively. The CSF obtained using stimulus-matched spatial pooling filters with no eye movements serves as the reference CSF (gray disks). All CSFs are computed for a 3 mm pupil, typical subject wavefront-based optics and eccentricity-based cone mosaics.

The CSFs derived using the SVM-Template-Energy computational observer in the absence and presence of fixational eye movements are depicted in Figure 3B by the red and blue disks, respectively. Note that the SVM-Template-Energy derived CSFs are nearly identical in the presence and absence of fixational eye movements, demonstrating that the sharp performance decline with spatial frequency can be eliminated when complex-cell like spatial energy mechanisms are used. This performance improvement at high spatial frequencies comes at a cost, however: an overall sensitivity drop by a factor of 2.0–2.5 across the entire frequency range independent of whether fixational eye movements are present or not (blue and red disks), as seen by comparison with performance of the SVM-Template-Linear observer in the absence of eye movements (gray disks).

### Impact of phototransduction

Next, we examined how phototransduction impacts contrast sensitivity. To isolate performance changes due to photo-transduction alone, we computed CSFs in the absence of fixational movements using the SVM-Template-Linear observer. The results of this analysis are depicted in Figure 4. The transformation in stimulus representation from cone excitations to photocurrents results in a contrast sensitivity loss of a factor of 2.0–2.5, with slightly more reduction at lower spatial frequencies. This spatial frequency effect is due is due to the increased down-regulation of photocurrent response gain at more eccentric retinal locations, where the cone excitation response is stronger due to the enlarged cone aperture diameters. More eccentric retinal locations come into play because the experiment we are modeling employed a fixed number of grating cycles, resulting in larger stimulus extents at lower spatial frequencies.

The sensitivity loss at the photocurrent processing stage will depend on stimulus conditions, particularly the mean luminance of the adapting background and also the stimulus duration. For example, as stimulus mean luminance increases we expect increased sensitivity loss at the level of photocurrent compared to excitations. This expectation is due to increased down regulation of photocurrent gain in the presence of a constant photocurrent noise for excitation rates up to 50,000-100,000 *R*^*^ × cone^−1^ × sec^−1 2^. Sensitivity will also depend on stimulus duration, due to temporal integration via the photocurrent temporal impulse response. Supplemental Figures 8 and 9 illustrate the effects of mean luminance and duration on the CSF for a range of spatial frequencies.

### Combined effect of fixational eye movements and phototransduction

Figure 5 depicts the combined effect of phototransduction and fixational eye movements. As we noted earlier, the use of the SVM-Template-Energy computational observer is effective at mitigating the effect of fixational eye movements, when applied at the level of cone excitations (Figure 3). When it is applied at the level of the photocurrent, it is less so. The reduced efficiency at the photocurrent stage occurs because the photocurrent impulse response temporally integrates the spatially-jitterred cone excitation response (see Figure 2F) *before* the energy mechanism has a chance to discount it.

None-the-less, spatial pooling via the energy mechanism is beneficial relatively to spatial pooling via a linear mechanism for maintaining performance at higher spatial frequencies. Better performance at higher spatial frequencies can be obtained by using ensembles of non-linear pooling mechanisms whose centers are spatially offset and which are tiling a region larger than the spatial extent of the stimulus (see Impact of spatial pooling mechanism in section Supplementary Material).

Overall, our results indicate that compared to the performance at the level of cone excitations without fixational eye movements, the combined effects of photocurrent encoding, fixational eye movements, and the energy based computational observer reduce performance by a factor of 5-10, in a manner that is largely but not completely independent of spatial frequency, with the spatial frequency dependence caused by the temporal integration of spatially jittered cone isomerization signals during phototransduction.

### Comparison of computational and human observer performance

To compare our computational-observer contrast sensitivity functions to human psychophysical sensitivity (Banks et al., 1987), we repeated the simulations for a 2 mm pupil to match the psychophysics. Figure 6 shows the comparison. The gray disks depict the CSF derived at the level of cone excitations in the absence of fixational eye movements using the SVM-Template-Linear observer (gray disks) and provide a reference CSF. The red disks depict performance at the level of photocurrent, also in the absence of fixational eye movements using the SVM-Template-Linear observer. The blue disks depict performance at the level of photocurrent in the presence of fixational eye movements, computed using the SVM-Template-Energy observer. Note that as spatial frequency increases, performance of the computational observer drops somewhat more rapidly than that of the human observer, so that the two CSFs are coming into agreement for spatial frequencies above about 20 c/deg. Thus, at the higher spatial frequencies, the early vision factors we model here account for the absolute contrast sensitivity of the human observers.

## Discussion

### Benefits and drawbacks of fixational eye movements

In the absence of fixational eye movements, post-receptoral processes render real human observers functionally blind to stationary objects (Riggs, Ratliff, Cornsweet, & Cornsweet, 1953), and recent studies have demonstrated that fixational eye movements can improve the precision of vision at intermediate (10 c/deg) spatial frequencies (Rucci, Iovin, Poletti, & Santini, 2007), and near the resolution limit (Ratnam, Domdei, Harmening, & Roorda, 2017). In contrast, our work suggests that fixational eye movements reduce sensitivity at high spatial frequencies. Because of this contrast, the relation between our work and theoretical work that has considered the role of fixational eye movements is worth discussion.

Rucci and colleagues (Rucci et al., 2007; Kuang, Poletti, Victor, & Rucci, 2012; Boi, Poletti, Victor, & Rucci, 2017), observed that fixational eye movements reformat the spatio-temporal power spectrum of the stimulus, and that under certain assumptions about post-receptoral processing this reformatting can be beneficial in terms of the transfer of information between the stimulus and subsequent post-receptoral mechanisms. Their theory is based on the observation that the temporal dynamics of ocular drift redistributes the 1*/f*^2^ power of static natural images by progressively boosting power at spatial frequencies up to 10-15 c/deg at nonzero temporal frequencies. This redistribution can improve detection performance for high spatial frequency targets if subsequent detection mechanisms are bandpass tuned to intermediate temporal frequencies (Boi et al., 2017). It can also ease the problem of decorrelating visual signals for efficient coding by capacity-limited post-receptoral mechanisms.

Our computational observer simulations, in contrast, do not show enhanced performance at high spatial frequencies in the presence of fixational drift, relative to performance when there are no fixational eye movements. Because of the apparent contradiction between our broad conclusion (fixational eye movements reduce performance at high spatial frequencies) and those of Rucci and colleagues (fixational eye movements improve performance at high spatial frequencies), it is worth highlighting differences between the two studies.

First, in our study, we explicitly model spatio-spectral low pass spatial filtering by the eye’s optics, isomerization noise, and the dynamics of phototransduction. These factors are not taken into consideration in the computation of the spatiotemporal power distribution by Rucci et al. (2007; Kuang et al., 2012; Boi et al., 2017), which is based on the pixel-level representation of the stimulus. Importantly, the low pass temporal filtering embodied in the conversion of cone excitations to photocurrent is not accounted for in their analysis. This filtering, when combined with fixational eye movements, leads to contrast reduction at high spatial frequencies, which post-receptoral processing cannot undo. Any advantage provided by reformatting of the spatio-temporal power spectrum must be large enough to overcome this loss.

Similarly, by starting their analysis with consideration of the power spectrum, the calculations of Rucci et al. (2007; Kuang et al., 2012; Boi et al., 2017), exclude temporal phase information. Our comparisons of the SVM-Template-Linear and SVM-Template-Energy observers show that removing phase information produces translation invariance in the face of fixational eye movements, but comes at a cost in sensitivity when applied in the case where there are no fixational eye movements. When considering possible functional benefits of fixational eye movements, the underlying cost of applying requisite translation-invariant decision models should also be taken into account.

There are other differences between the two studies. We model a 100 msec stimulus as we are interested in comparing with the data of Banks et al. (1987). During this short time there is not a significant reduction in the photocurrrent response amplitude (see Figure 11E) that drift transients could enhance. Boi et al. (2017) report that contrast sensitivity enhancement at high spatial frequencies in the presence of fixational drift is nearly absent for stimuli presented for 100 msec and that a progressive enhancement occurs during prolonged stimulus exposure (800 msec). Also, in our simulations, stimuli at progressively higher spatial frequencies have progressively reduced spatial extent, maintaining a constant number of cycles. In the study of Rucci and colleagues (Rucci et al., 2007; Kuang et al., 2012; Boi et al., 2017), stimuli of high and low spatial frequencies were matched in size. This may cause fixational eye movements to have a larger effect for our high spatial frequency stimuli than for the stimuli employed by Rucci and colleagues.

**Figure 8:**
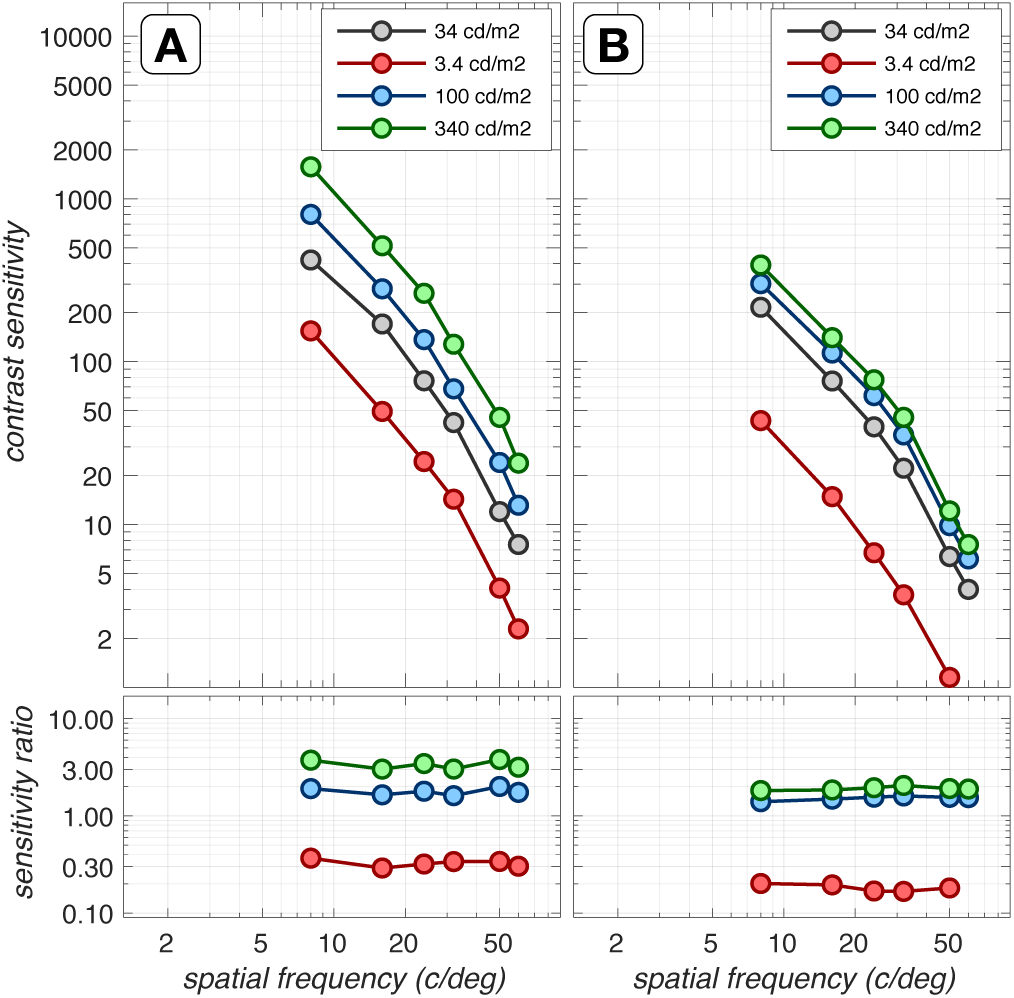
Impact of background luminance [Suppl. Fig. S1]. Contrast sensitivity functions computed at different background levels, based on the isomerization (left) and the photocurrent responses (right). These CSFs were computed for a 3 mm pupil, typical subject wavefront-based optics, eccentricity-based cone mosaics, and in the absence of eye movements using a linear computational observer. The reference CSF (gray disks) is computed for 34 cd/m^2^, the background used in all computations in this paper. At the level of isomerizations (**A**), the ratios of sensitivity with respect to the reference CSF follows the square root law of Poisson noise limited sensitivity. At the level of photocurrents (**B**), where noise is additive and sensitivity is regulated by the phototransduction dynamics, the contrast sensitivity increase with increasing luminance is closer to Weber’s law which implies no change in contrast sensitivity with luminance. Note that our photocurrent model has been validated against experimental data up to a luminance of about 100 cd/m^2^, so the performance shown for 340 cd/m^2^ represents an extrapolation.

**Figure 9:**
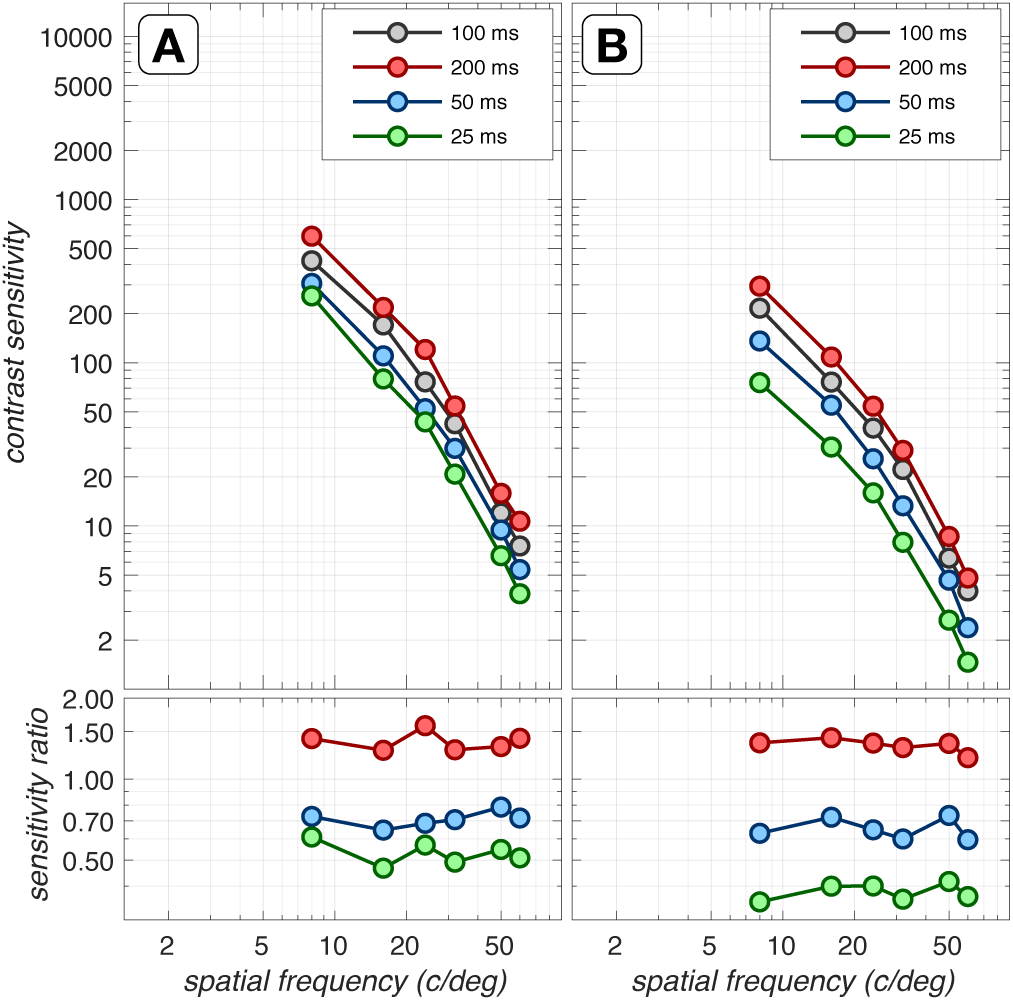
Impact of stimulus duration [Suppl. Fig. S2]. Contrast sensitivity functions computed for different stimulus durations, at the level of isomerizations (left) and photocurrents (right). These CSFs were computed for a 3 mm pupil, typical subject wavefront-based optics, eccentricity-based cone mosaics, and in the absence of eye movements using a linear computational observer. The reference CSF (gray disks) is computed for 100 msec, the duration used in all computations in this paper. Note that at the level of isomerizations, ratios of CSFs with respect to the reference CSF cluster around 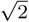 for 200 msec, 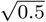 for 50 msec, and 0.5 for 25 msec, as expected from the square root law of Poisson noise limited sensitivity. At the level of photocurrents, where noise is additive and sensitivity is regulated by the phototransduction dynamics, there is a more dramatic effect of stimulus duration on performance.

**Figure 10:**
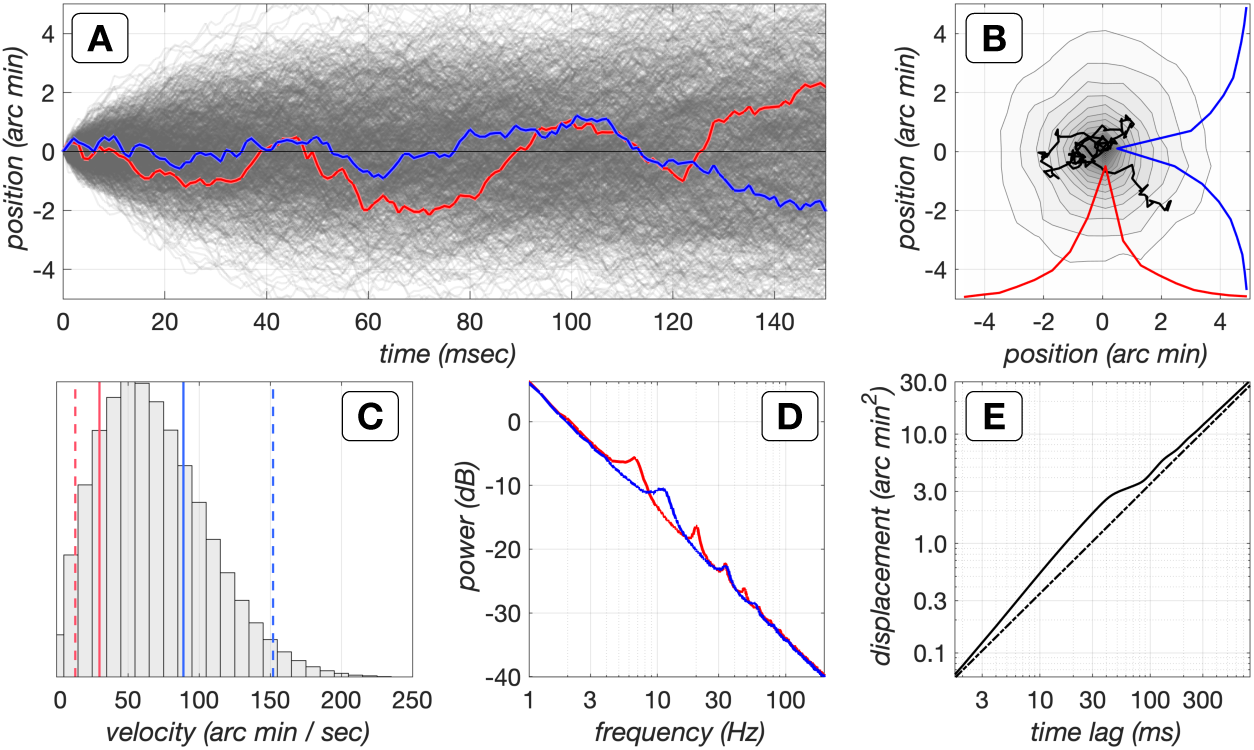
Dynamics of drift fixational eye movements generated by our model. **A.** X-/Y-trajectories of 1024 eye movement paths during a period of 150 msec. For visualization purposes, the paths always start at (0,0), but in the simulations each path starts at a random location and the centroid of each path is constrained to be at (0,0). Red and blue lines denote the x/y trajectories of a single eye movement path. **B**. Fixation span of the 1024, 150 msec long eye movement paths. The black line depicts the eye movement path whose x/y trajectories are depicted in **A. C.** Distribution of instantaneous velocities in 1024 model eye movement paths. The red and blue solid lines depict the mean drift velocities from 2 human subjects that had the lowest and highest velocity, respectively, in a pool of 12 subjects (Cherici et al., 2012). The red and blue dashed lines represent *−*1*σ* and +1*σ* of the drift velocity distributions for these subjects, respectively. **D.** Power spectra of the x and y-drift trajectories. The spectral peaks are due to the oscillatory behaviour of the modeled delayed negative feedback mechanism, which has slightly different delays for the x- and y-drift components. **E.** Displacement analysis of drift eye movement paths. The solid line depicts the mean squared displacement, 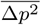, as a function of time lag, for the generated eye movement paths. The dashed line depicts 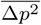for a purely Brownian process. Notice the persistent behaviour between 2 to 30 msec, which causes the eye to diffuse more than what it would have if it were under the control of a purely Brownian process, and the antipersistent behaviour which stabilizes eye position at longer time delays (100 to 500 msec).

**Figure 11:**
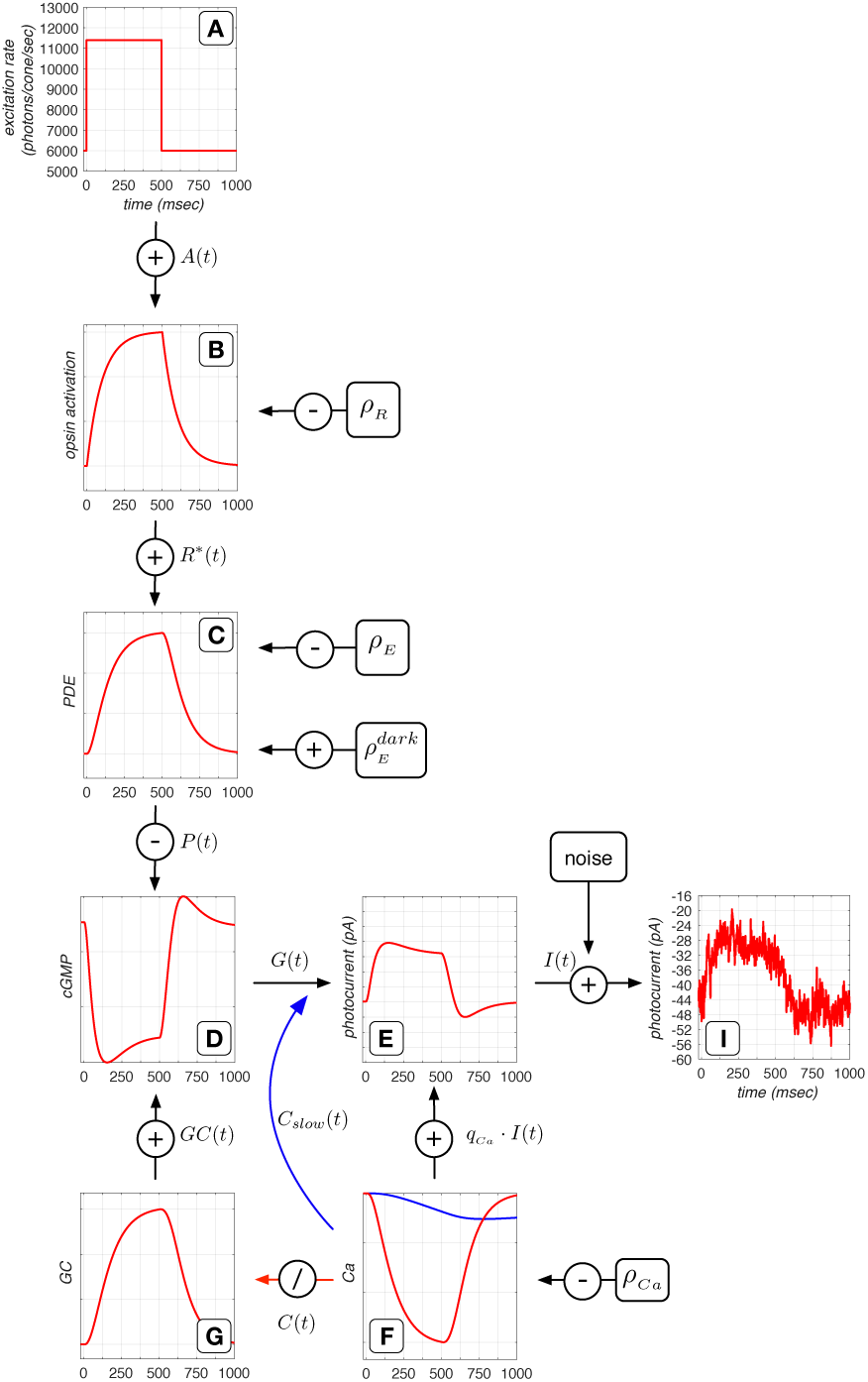
Phototransduction cascade model. Depicted here are the responses of the different model components to a 500 msec light increment pulse. **A.** Sequence of mean cone excitation rate. **B.** Opsin activation in response to the change in cone excitation rate. **C.** Activation of PDE enzymes in response to opsin activation. **D.** Change in cGMP concentration, which is synthesized by CG and broken down by PDE enzymes. **E.** Membrane current, photocurrent, which is an instantaneous function of the cGMP activation. **F.** Change in intracellular Ca^+2^ concentration, which is the result of inflow via the membrane current and outflow, via the Na^+^- Ca^+2^ exchanger pump. Two feedback mechanisms both based on the intracellular Ca^+2^ concentration modify the membrane current. A slow Ca^+2^-derived signal which directly regulates the membrane current, and an indirect signal which modulates production of GC, shown in **G.**, which is responsible for producing cGMP. See text for more details. **I.** Noisy membrane current response instance generated by adding photocurrent noise to the mean membrane current response (**E**).

In the other direction, our present model does not consider limits imposed by known post-receptoral mechanisms. These include both the spatio-temporal filtering applied by circuits in the retina, as well as bandwidth limits on the transmission of information between the retina and the cortex. The work of Rucci and colleagues, on the other hand, focuses on the relation between these factors (e.g., spatiotemporal filtering by ganglion and cortical neurons) and the effect of fixational eye movements. We are planning to extend our modeling to explicitly include post-receptoral mechanisms (e.g. the properties of retinal ganglion cells), at which point it will be of interest to examine again the effects of fixational eye movements. Modeling both the factors we consider here and those underlying the thinking of Rucci and colleagues seems likely to further clarify the costs and benefits of fixational eye movements and how they depend on what is taken as given about post-receptoral processing.

### More sophisticated inference engines

Another theoretical approach to understanding the role of fixational eye movements in vision was developed by Anderson, Olshausen, Ratnam, and Roorda (2016). Their work considers inference engines that seek to simultaneously estimate both the visual stimulus and the eye movement path, thus minimizing the effects of spatial uncertainty introduced by the eye movements. They show potential advantages of fixational eye movements, particularly if cone sampling density is low relative to the spatial structure in the stimulus. In that case, the sweep of the mosaic across the retinal image samples stimuli more finely that a stationary retina. It is thus possible that introducing inference engines of this sort may lead to a lower performance loss, or even a performance gain, in the face of fixational eye movements. Such inference engines might also help explain the stability of our perceptual representations in the face of fixational eye movements. Note, however, that inference engines of the sort described by Anderson et al. (2016) require some short-term storage and cannot be implemented before the cone-bipolar synapse; they remain subject to information loss caused by the temporal smearing of cone excitation responses due to photocurrent encoding (Figure 2).

More generally, other inference engines, for example those based on deep convolutional networks, may have a lower performance loss than the SVM-Template based inference engines we studied. Also, it is possible that performance loss may be reduced when multiple inference mechanisms are employed. For example, at low to mid spatial frequencies (2-8 c/deg), where retinal jitter due to eye movements causes essentially zero performance degradation when a linear pooling based inference engine is used, it would be detrimental to use energy based inference engines which have reduced overall sensitivity. Further, at very high spatial frequencies it would be beneficial to use ensembles of energy based mechanisms as we demonstrate in the supplementary section Impact of spatial pooling mechanism. Hybrid inference engines that rely more on the outputs of linear summation mechanisms for stimuli of low to mid spatial frequencies and more on the outputs nonlinear summation mechanisms for stimuli of high spatial frequencies should also be examined. An important direction for future work is to couple the study of sophisticated inference engines with model visual systems such as the one we present here, that incorporate the known limits of visual encoding.

### Caveats

We list some caveats with respect to the completeness of our modeling of early vision. First, we do not account at a fine scale for eccentricity-based changes in photocurrent sensitivity. In our present simulations, the amplitude and dynamics of the light-regulated photocurrent impulse response are determined based on the average activation of each cone type across the entire mosaic. This approximation improves computational efficiency and affects our simulations in two ways. Central cones, which have the smallest apertures, have lower excitation levels than the mean excitation level across the entire mosaic, and therefore, for these cones, the modeled photocurrent impulse response is more heavily regulated than what it would have been if the actual cone excitation was used. Since the model photocurrent noise is stimulus independent, the signal to noise ratio for central cones is slightly lower that what it ought to be. The opposite holds for peripheral cones: their photocurrent responses have higher signal to noise ratio than they ought. Also, foveal cones are considerably slower than peripheral cones (Sinha et al., 2017), so the temporal integration of cone excitation signals in the presence of fixational eye movements would have less of an impact for peripheral stimulus locations than what is captured by our simulations, which assume foveal cone photocurrent dynamics at all mosaic locations. The combined effect probably results in a lower overall sensitivity, and we suspect that our CSF estimates would be a little higher if we took eccentricity-based changes in photocurrent sensitivity and dynamics into account.

Second, our simulations employed stimulus-independent photocurrent noise. Work by Angueyra and Rieke (2013) has shown that noise is not entirely independent of the background cone excitation level. Low frequency components (1-10 Hz) are subject to the same adaptational gain reduction as the mean photocurrent response, whereas high-frequency components (> 100 Hz) are less affected by background light level. So in real cone mosaics, in which peripheral cones have larger apertures than central cones, the noise spectrum changes shape with eccentricity. Our simulations do not capture this effect.

Third, the simulations match the size of the cone mosaic to the stimulus size, which in turn varies with spatial frequency. This simplification was chosen for computational efficiency. However, this poses a problem for the highest spatial frequency stimuli, for which part of the retinal image can be brought out of the field of view of the cone mosaic in the presence of fixational eye movements. The supplementary section Impact of spatial pooling mechanism presents an analysis of performance for different spatial pooling mechanisms which extend beyond the stimulus spatial support.

Fourth, our fixational eye movement model computes drift eye movement trajectories whose mean velocity is 60 arc min/sec (Figure 10C), near the mean of the velocity distribution across a number of human observers (Cherici et al., 2012). However, Cherici et al. (2012) report that trained observers have significantly lower mean drift velocities (30 arc min /sec) as opposed to naive observers, who can have mean drift velocities up to 90 arc min/sec, and therefore trained observers have narrower fixation spans. The observers employed by Banks et al. (1987) were the authors themselves, and were certainly well trained. Our computational observer performance would likely improve in the higher spatial frequency range if we employed fixational eye movements whose velocity matched the low end of the distribution of velocities reported by Cherici et al. (2012).

## Summary and conclusion

We extended the ISETBio computational observer model of the human spatial contrast sensitivity (CSF) to incorporate fixational eye movements and the transformation of quantal cone excitations to photocurrent. An introductory script that demonstrates computation of photocurrent responses in the presence of fixational eye movements can be found at https://github.com/isetbio/ISETBioCSF/tree/master/tutorials/recipes/CSFpaper2.

Our results indicate that fixational eye movements abolish sensitivity above 10 c/deg for a computational observer that employs a stimulus-matched, linear pooling template. Energy based computational observers eliminate the sharp performance decline at high spatial frequencies, but at the cost of a decrease in sensitivity. The decrease in overall sensitivity in the absence of eye movements should not be surprising – to achieve translation-invariance, energy-based observers ignore the stimulus spatial phase, using less stimulus information than the linear summation observer.

Phototransduction-induced sensitivity regulation and additive noise also decreases sensitivity by a factor of 2. Combining the effects of fixational eye movements, photocurrent encoding, and energy-based computational observer, brings the computational observer performance to levels that are within a factor 1 to 2 of human sensitivity, depending on spatial frequency. This analysis indicates that the sensitivity loss observed in human performance relative to the sensitivity of an ideal observer at the cone excitations can be largely be accounted for by fixational eye movements, photocurrent encoding, and the performance of the inference engines we considered. This leaves little room for additional sensitivity loss as the signal is processed by the very large number of neurons in thalamus and cortex.

## Supplementary Material

### Impact of spatial pooling mechanism

In the constant cycles paradigm, as stimulus spatial frequency increases stimulus size becomes progressively smaller and so does the spatial extent of the spatial pooling mechanism, which is matched to stimulus area. Fixational eye movements can move the retinal image of the stimulus outside the field of view of these pooling mechanisms, thereby lowering performance.

Here we examine how performance is affected by choosing spatial pooling regions that extend beyond the stimulus. We consider two spatially-extended cone pooling schemes. The first scheme, depicted in Figure 7B consists of a single spatial pooling energy mechanism which is centered on the retinal image but which spans a spatial region larger than the stimulus. The quadrature-phase pair pooling kernels of this energy mechanism are depicted in Figures 7D and 7E. The second scheme, depicted in Figure 7C, consists of an ensemble of spatial pooling energy mechanisms whose centers are positioned on a spatial grid which extends beyond the stimulus spatial support. The example shown in Figure 7C is for a 3×3 grid. For this mechanism the input to the SVM classifier is the ensemble of temporal responses of 9 spatial filters. The cos-phase pooling kernels for 2 of these mechanisms (outlined in red and orange in Figure 7C are depicted in Figures 7F and 7G, respectively.

The high spatial frequency CSF regimes computed using different spatial pooling mechanisms are depicted in Figure 7H for cone isomerization signals, and Figure 7I for photocurrent signals. The CSFs computed using the stimulus-matched pooling mechanism in the absence of fixational eye movements is depicted by the gray disks and serve as reference CSFs. The remaining CSFs are computed in the presence of fixational eye movements. Red disks depict the CSFs derived using the stimulus-matched energy template, whereas the green and blue disks depict the CSFs derived using a single energy filter that extended beyond the spatial support of these high frequency stimuli, 0.33 degs and 0.5 degs, respectively. Finally, the CSFs derived using ensembles of spatially-offset energy filters are depicted in magenta (0.3 degs, 3×3 grid) and orange (0.5 degs, 5×5 grid).

Note that at the level of isomerizations (Figure 7H), the single spatially-extended pooling mechanisms (0.3 and 0.5 degs) both perform worse than the stimulus-matched spatial pooling mechanism. On the other hand, the mechanisms consisting of a 3×3 and a 5×5 ensemble of spatial pooling filters significantly outperform the stimulus-matched spatial pooling mechanism at the highest spatial frequencies (40, 50 and 60 c/deg), even in the absence of fixational eye movements (gray disks). At the level of photocurrents (Figure 7I), the ensemble of spatial pooling filters mechanisms again outperform both the stimulus-matched and the spatially-extended single spatial pooling mechanism at 40 and 50 c/deg, whereas at 60 c/deg we were not able to obtain a detection threshold for any spatial pooling mechanism.

Overall, these results indicate that in the presence of fixational eye movements, ensembles of spatially-overlapping energy mechanisms can enhance performance at high spatial frequencies whereas spatially extended and stimulus matched mechanisms cannot. Performance of the ensemble mechanisms depends on the amount of spatial overlap and the width of the component filters, but this dependence is not examined in the present work.

**Impact of background luminance**

**Impact of stimulus duration**

## Methods

### Modeling optics and cone mosaic excitation

ISETBio computations begin with a quantitative description of the visual stimulus, here, a spatio-temporal pattern specified as the spectral radiance emitted at each location and time on a flat screen. The spectral irradiance incident at the retina (retinal image) is computed by taking into account human optical factors such as pupil size, wavelength-dependent blur, on-axis wavefront aberrations, and wavelength-dependent transmission through the crystalline lens. Cone mosaic excitations are computed from the spectral retinal irradiance using naturalistic cone mosaics which model the relative number of L, M and S cones, the existence and size of an S–cone free zone in the central fovea, the variation in cone spacing, inner segment aperture size and outer segment length with eccentricity, cone photopigment density, and macular pigment density. Performance in a two-alternative forced choice detection task is assessed using support-vector-machine based binary classifiers which operate on the output of mechanisms which pool cone responses over space using linear and non-linear (energy-based) stimulus-derived pooling schemes. Detailed descriptions of all elements of this computational pipeline and estimates of how different elements impact the spatial contrast sensitivity function can be found in Cottaris et al. (2019). In the following sections, we describe the ISETBio modeling of two additional elements of early vision, fixational eye movements and phototransduction, whose impact on the spatial contrast sensitivity function is examined in the present paper.

### Modeling fixational eye movements

The fixational eye movement model in ISETBio includes a drift and a microsaccade component. The drift component is generated by a delayed feedback loop mechanism based on work by Mergenthaler and Engbert (2007), which generates eye movement paths using a generalized Brownian motion process. The microsaccade component injects abrupt shifts in eye position with the onset, speed, and amplitude of the trajectories drawn from published distributions of microsaccades (Martinez-Conde et al., 2009). Saccade direction is based on heuristics which aim to maintain fixation while also avoiding recently visited positions. In the present work, which simulates presentation of a 100 msec stimulus, we only engage the drift component, as microsaccades typically occur only every 500 to 1000 msec (Martinez-Conde et al., 2009).

#### Drift fixational eye movement generation model

Drift eye movement trajectories during fixation of steady targets resemble generalized Brownian motion (Engbert & Kliegl, 2004). In a generalized Brownian motion process, the mean square displacement, 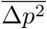, at a time lag relative to an arbitrary time point, Δ*T*, computed as 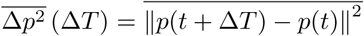, is proportional to Δ*T*^*K*^, where *K* is a real number between 0 and 2. In a purely Brownian process, the position at time step *i* + 1, *p*_*i*+1_, is given by

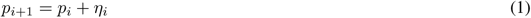

where *p*_*i*_ is the position at time step *i*, and *η*_*i*_ is a normally distributed random variable with zero mean. In such a process, the sequence of spatial displacements is uncorrelated and *K* = 1. Using diffusion analysis, Engbert and Kliegl (2004) showed that fixational eye movements differ from pure Brownian motion, exhibiting correlations over two time scales. Over short time scales (2 to 30 msec), the eye has a tendency to continue to move in the current direction (persistent behavior), resulting in correlated displacements and *K* > 1. Over longer time scales (100 to 500 msec), there is a tendency to reverse direction (anti-persistent behavior), resulting in uncorrelated displacements and *K* < 1.

Differences in eye position dynamics from those of a purely Brownian process affect fixation span and may affect performance. To simulate a combination of persistent and anti-persistent dynamics, we generate fixational eye movements using the delayed random walk model proposed by Mergenthaler and Engbert (2007). In this model the position at time step *i* + 1, *p*_*i*+1_, is given by

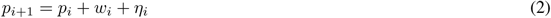

where

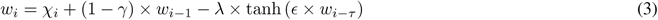

In equation 3, the autoregressive term, (1 − *γ*) × *w*_*i*−1_, generates the persistent behavior at short time scales. The negative delayed feedback term, −λ × tanh (*ϵ* × *w*_*i*−*τ*_), generates the anti-persistent behaviour at longer time scales. *χ*_*i*_ represents a Gaussian random noise process with zero mean, which provides the driving signal for *w*_*i*_. The values of the model parameters that we used in the present study are listed in Table 1 and are taken from Mergenthaler and Engbert (2007). The model generates fixational eye movement paths with a resolution of 1 msec, with X- and Y-position coordinates computed independently.

**Table 1:** Values of the drift fixational eye movement model parameters used in this study.

| Symbol | Description | Value |
| --- | --- | --- |
| $\gamma$ | control gain | 0.25 |
| $\mu_\chi$ | control noise $\chi$, mean | 0.000 |
| $\sigma_\chi$ | control noise $\chi$, st.dev. | 0.075 |
| $\mu_\eta$ | position noise $\eta$, mean | 0.000 |
| $\sigma_\eta$ | position noise $\eta$, st.dev. | 0.35 |
| $\lambda$ | feedback gain | 0.15 |
| $\epsilon$ | feedback steepness | 1.1 |
| $\tau_x$ | feedback delay (x-position) | 0.07 |
| $\tau_y$ | feedback delay (y-position) | 0.04 |

#### Dynamics of model drift fixational eye movements

Various properties of the eye movement paths generated by the model are illustrated in Figure 10. Figure 10A illustrates the X-/Y-position components of 1024 drift trajectories, each lasting for 150 msec. Figure 10B depicts the fixation span computed over all traces during this duration, which is defined as the spatial probability distribution with which the eye is at each position during the analyzed period. The distribution of instantaneous drift velocities, computed by taking the time derivative of the low passed position signal (low pass filter: 41-sample Savitzky-Golay filter), is depicted in Figure 10C. Notice that these model velocities span the range of velocities measured in human observers (Cherici et al., 2012). The X- and Y-position spectra are displayed in Figure 10D, in red and blue lines, respectively. The displacement of eye position as a function of time lapsed from any time point during the trajectory is depicted by the solid line in Figure 10E. Note that for short time scales (2 to 30 msec), displacement is larger from what would be observed if the motion were purely Brownian (dashed line), whereas for longer time scales (100-500 msec), anti-persistent dynamics minimize this difference.

### Modeling photocurrent generation

Our model of cone photocurrent response consists of two stages. In the first stage, a biophysically-based model of the phototransduction cascade transforms a time-varying sequence of photon excitation rate to a photocurrent temporal response. The model is a modified version of the canonical model of phototransduction (Pugh & Lamb, 1993; Hateren, 2005) with parameters based on population recordings from primate cone photoreceptors (Angueyra, 2014). Although we have implemented the full non-linear conversion between time-varying excitation rate and photocurrent, the full calculation is compute-intensive because a small time step (0.1 msec) is required in order to accurately simulate the differential equations that govern the phototransduction cascade. Here we use the full calculation to determine a linear approximation which is valid for near-threshold perturbations around a mean luminance. That is, the full phototransduction model is used to compute a photocurrent impulse response function, which is defined as the outer segment membrane current in response to a cone excitation delta function superimposed on a constant cone excitation background rate. This biophysically-derived impulse response function is specific to the stimulus mean luminance, and stimuli of different mean luminances and chromaticities will have different photocurrent impulse responses. The derived photocurrent impulse response is downsampled to the time step of the simulations, here 5 msec, and subsequently convolved with the sequence of the mean cone excitations, which are also computed every 5 msec, to derive the noise-free photocurrent response.

The second stage of the model adds a stochastic component which captures noise in the phototransduction cascade. The noise has Gaussian amplitude distribution and a power spectrum that is matched to that of primate cone photoreceptors (Angueyra & Rieke, 2013). The computed photocurrent captures the characteristics of primate cone responses (Angueyra, 2014) for a range of adaptation levels, 0 to 30,000 R^*^ × cone^−1^ × sec^−1^. In this range, photopigment bleaching is less than 2%, assuming a half-bleaching constant of 6.4 log R^*^ × cone^−1^ × sec^−1^, and can therefore be ignored. The half-bleaching constant was estimated from the value of 4.3 log Trolands provided by Rushton and Henry (1968).

#### Biophysically-based model of the phototransduction cascade

In darkness, there is a constant inflow of Na^+^ and Ca^+2^ ions into the photoreceptor outer segment via cyclic guanosine monophosphate (cGMP)-gated channels, many of which are open due to the high concentration of intracellular cGMP. cGMP is constantly being produced by the enzyme guanylate cyclase (GC). The constant inflow of Na^+^ and Ca^+2^ ions into the photoreceptor creates a negative current. This negative current hyperpolarizes the cone membrane, which results in a continuous release of glutamate at the synapses with bipolar and horizontal cells. When a photon isomerizes an opsin molecule, it initiates a biochemical cascade which results in the activation of multiple phosphodiesterase (PDE) enzymes. The increased PDE activity hydrolizes cGMP at a higher rate than in the dark, thereby reducing the intracellular cGMP concentration, which leads to closure of cGMP-channels. This blocks the entry of Na^+^ and Ca^+2^ ions into the cone, causing a depolarization in the membrane and a decrease in glutamate released. Our model of this process, which captures the steps between isomerization and the modulation of membrane current, is illustrated in Figure 11. The implementation of the different stages is as follows.

##### Opsin activation

Absorption of photons by photopigment molecules turns inactive opsin proteins, *R*, into their activated state, *R*^*^. Activated opsin molecules are produced instantaneously with a rate that is proportional to the photon absorption rate, *A*(*t*), and inactivated with a rate constant, *ρ* _*R*_. When the light intensity is such that *A*(*t*) < 30, 000 photons/cone/sec, we can neglect photopigment bleaching and treat the concentration of inactive photopigment as a constant. In this regime, the production of activated opsin, *R*^*^(*t*), is described by

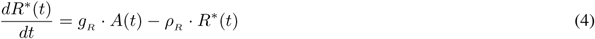

where *g*_*R*_ is a scaling constant.

##### PDE concentration

Phosphodiesterase (PDE) enzymes are in turn activated by activated opsin proteins with a rate 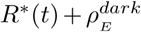, where 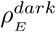, is the spontaneous PDE activation rate in the dark, and become inactivated with a rate constant, *ρ* _*E*_. The production of activated PDE, *E*(*t*), is described by:

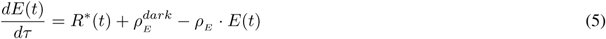

##### cGMP concentration

cGMP molecules are synthesized continuously due to the GC protein activity at a rate *GC*(*t*), and hydrolized at a rate that is product of PDE enzymatic activity, *P*(*t*), and cGMP concentration, *G*(*t*). The concentration of cGMP, *G*(*t*), is described by:

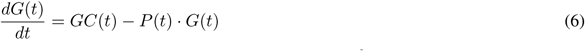

The rate at which GC is producing cGMP is an instantaneous function of the intracellular Ca^+2^ concentration, *C*(*t*):

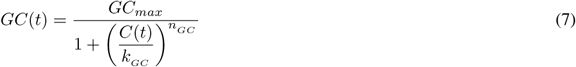

where *GC*_*max*_ is the maximal production rate, *k*_*GC*_ is the half-maximal Ca^+2^ concentration and *n*_*GC*_ is an exponent that determines the steepness of the relation between *GC*(*t*) and *C*(*t*).

##### *Ca*^+2^ *concentration*

The intracellular Ca^+2^ concentration, *C*(*t*), depends on two factors: the Ca^+2^ inflow though the open cGMP-gated outer segment membrane channels and the Ca^+2^ outflow via Na^+^– Ca^+2^ exchanger pumps, and is described by:

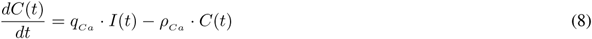

where 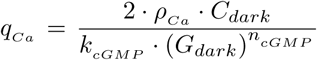 is the fraction of the ionic membrane current that is carried by Ca^+2^ ions, and *ρ* _*Ca*_ is the rate constant at which Na^+^ – Ca^+2^ exchanger pumps eject Ca^+2^ out of the receptor. Note that Ca^+2^ drives a negative feedback pathway in the outer segment, since the concentration of Ca^+2^ regulates the rate at which cGMP is produced by GC (equation 7).

##### Photocurrent

The photocurrent, *I*(*t*), is the ionic inflow of extracellular Na^+^ and Ca^+2^ into the photoreceptor outer segment. Na^+^ and Ca^+2^ enter via pores located in the outer segment plasma membrane which remain open when cGMP molecules bind to them. The number of open cGMP-gated channels depends on the cGMP concentration, *G*(*t*), and determines the amplitude of the photocurrent. In the canonical phototransduction model, *I*(*t*) is an instantaneous function of *G*(*t*), and is described by:

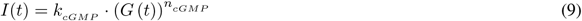

where *k*_*cGMP*_ and *n*_*cGMP*_ are constants that determine the non-linear dependence of current on open cGMP channels. In our model, *I*(*t*) is regulated based on a slow Ca^+^2 – derived signal, *C*_*slow*_(*t*), as follows:

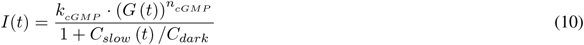

where *C*_*dark*_ is the Ca^+2^ concentration in the dark. The *C*_*slow*_(*t*) signal tracks the calcium concentration, *C*(*t*), filtered through a slow rate constant, 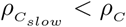, and is described by:

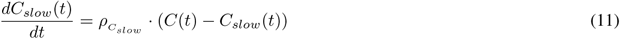

This is a second Ca^+2^ – based feedback pathway in the outer segment which provides a slow adaptational mechanism that helps to capture cone responses to impulse, step, and naturalistic stimuli in the primate (Angueyra, 2014). The values of all parameters of this cone photocurrent model are listed in Table 2.

**Table 2:**
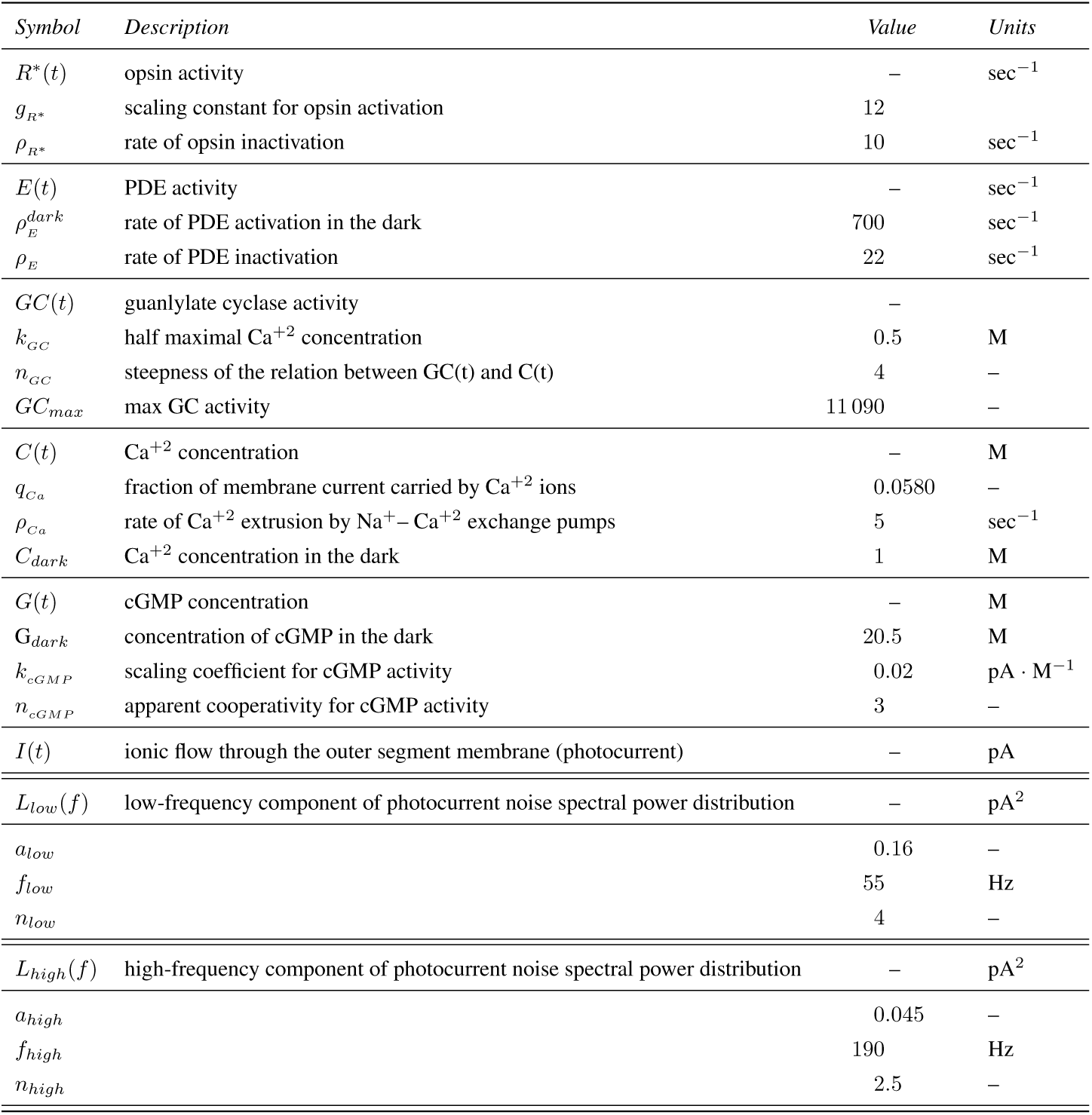
Parameters of the cone photocurrent model used in the present study. Parameter values were determined by fitting the model to primate cone responses to impulses delivered in darkness, pulses on various adapting backgrounds, and naturalistic stimuli (Angueyra, 2014).

| Symbol | Description | Value | Units |
| --- | --- | --- | --- |
| $R^*(t)$ | opsin activity | – | $\text{sec}^{-1}$ |
| $g_{R^*}$ | scaling constant for opsin activation | 12 | |
| $\rho_{R^*}$ | rate of opsin inactivation | 10 | $\text{sec}^{-1}$ |
| $E(t)$ | PDE activity | – | $\text{sec}^{-1}$ |
| $\rho_E^{dark}$ | rate of PDE activation in the dark | 700 | $\text{sec}^{-1}$ |
| $\rho_E$ | rate of PDE inactivation | 22 | $\text{sec}^{-1}$ |
| $GC(t)$ | guanylate cyclase activity | – | |
| $k_{GC}$ | half maximal $\text{Ca}^{+2}$ concentration | 0.5 | M |
| $n_{GC}$ | steepness of the relation between GC(t) and C(t) | 4 | – |
| $GC_{max}$ | max GC activity | 11 090 | – |
| $C(t)$ | $\text{Ca}^{+2}$ concentration | – | M |
| $q_{Ca}$ | fraction of membrane current carried by $\text{Ca}^{+2}$ ions | 0.0580 | – |
| $\rho_{Ca}$ | rate of $\text{Ca}^{+2}$ extrusion by $\text{Na}^{+}$– $\text{Ca}^{+2}$ exchange pumps | 5 | $\text{sec}^{-1}$ |
| $C_{dark}$ | $\text{Ca}^{+2}$ concentration in the dark | 1 | M |
| $G(t)$ | cGMP concentration | – | M |
| $G_{dark}$ | concentration of cGMP in the dark | 20.5 | M |
| $k_{cGMP}$ | scaling coefficient for cGMP activity | 0.02 | $\text{pA} \cdot \text{M}^{-1}$ |
| $n_{cGMP}$ | apparent cooperativity for cGMP activity | 3 | – |
| $I(t)$ | ionic flow through the outer segment membrane (photocurrent) | – | pA |
| $L_{low}(f)$ | low-frequency component of photocurrent noise spectral power distribution | – | $\text{pA}^2$ |
| $a_{low}$ | | 0.16 | – |
| $f_{low}$ | | 55 | Hz |
| $n_{low}$ | | 4 | – |
| $L_{high}(f)$ | high-frequency component of photocurrent noise spectral power distribution | – | $\text{pA}^2$ |
| $a_{high}$ | | 0.045 | – |
| $f_{high}$ | | 190 | Hz |
| $n_{high}$ | | 2.5 | – |

#### Photocurrent noise model

Photocurrent noise is generated by multiplying the Fourier transform of Gaussian random noise with the sum of two spectral functions, *L*_*low*_(*f*) and *L*_*high*_(*f*), and subsequently computing an inverse Fourier transform. The *L*_*low*_(*f*) and *L*_*high*_(*f*) functions are given by

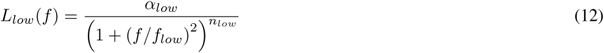

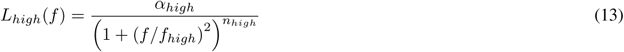

and model the low and high temporal frequency components, respectively, of the spectral power distribution of photocurrent noise recorded in macaque cone photocurrent responses. The values of the parameters of *L*_*low*_(*f*) and *L*_*high*_(*f*) are listed in Table 2. Figure 12A depicts the spectral power distribution of the generated noise (black line) along with and, and Figure 12B depicts 150 msec of the generated noise, along with a histogram of the amplitude distribution accumulated over 1024 instances.

**Figure 12:**
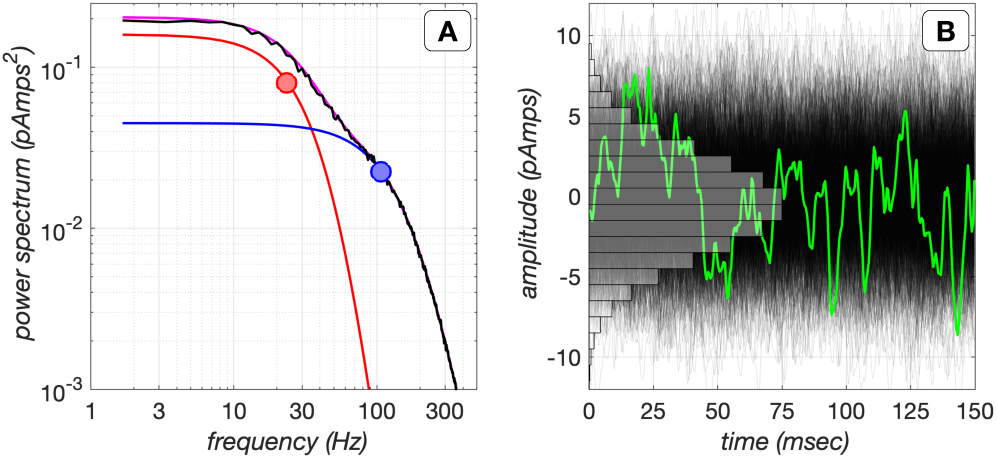
Photocurrent noise model. **A.** Photocurrent noise is generated by shaping the spectral power distribution of Gaussian noise to that corresponding to the sum of two spectral functions, and depicted in red and blue lines, with corner frequencies of 23 Hz and 107 Hz, respectively. Their sum is depicted in magenta, and the black line depicts the spectral power distribution of the realized noise. **B.** 1024 instances of 150 msec long photocurrent noise generated by the model. The green line depicts a single noise instance. The amplitude histogram of the noise is depicted in gray.

Angueyra and Rieke (2013) have shown that the low frequency noise component includes contributions from extrinsic noise (noise due to fluctuations in cGMP concentration due to spontaneous activation of Opsin and PDE) and intrinsic noise (noise in the opening and closing of cGMP channels), whereas the high frequency noise component is mostly due to intrinsic noise. Moreover, Angueyra and Rieke (2013) have shown that the power spectrum of the low frequency noise component decreases with background levels (similarly to the gain of mean photocurrent response) and has a half-desensitizing level of 4500 *R*^*^ × cone^−1^ × sec^−1^, whereas the power spectrum of the high frequency component is mildly affected by the background, with a shallow rate that is not proportional to the inverse of the background, with a half-desensitizing level of 17500 *R*^*^ × cone^−1^× sec^−1^. These background-dependent noise gain effects are not included in our implementation, in which the amplitude of the high and low frequency components are fixed.

## Acknowledgments

Supported by the Simons Foundation Collaboration on the Global Brain Grant 324759 and Facebook Reality Labs.

## Footnotes

1 In the ISETBio software, the term more general term optical image is used to refer to the retinal image. In this paper, however, we will use the term retinal image.

2 As cone excitation rates exceed 50,000-100,000 *R*\* × cone^−1^ × sec^−1^, photocurrent noise starts to decrease with background level (Angueyra, 2014). This decrease is not implemented in our present photocurrent model, which implements a constant amplitude of photocurrent noise.

